# Human intestinal tissue-resident memory CD8+ T cells comprise transcriptionally and functionally distinct subsets

**DOI:** 10.1101/869917

**Authors:** Michael E.B. FitzPatrick, Nicholas M. Provine, Lucy C. Garner, Kate Powell, Ali Amini, Sophie Irwin, Helen Ferry, Tim Ambrose, Peter Friend, Georgios Vrakas, Srikanth Reddy, Elizabeth Soilleux, Paul Klenerman, Philip J. Allan

**Affiliations:** Translational Gastroenterology Unit, Nuffield Department of Medicine, University of Oxford, Oxford, UK; Peter Medawar Building for Pathogen Research, University of Oxford, UK; Oxford Transplant Centre, Oxford University Hospitals NHS Foundation Trust, Churchill Hospital, Oxford, UK; Division of Cellular and Molecular Pathology, Addenbrooke’s Hospital, Cambridge, UK; NIHR Biomedical Research Centre, John Radcliffe Hospital, Oxford, UK

## Abstract

Tissue-resident memory T (T_RM_) cells provide key adaptive immune responses in infection, cancer, and autoimmunity. However transcriptional heterogeneity of human intestinal T_RM_ cells remains undefined, and definitive markers of CD103-T_RM_ cells are lacking. Here, we investigated transcriptional and functional heterogeneity of human T_RM_ cells through the study of donor-derived intestinal T_RM_ cells from intestinal transplant recipients. Single-cell transcriptional profiling identified four conventional T_RM_ populations, with two distinct transcriptional states of CD8+ T_RM_ cells, delineated by *ITGAE* and *ITGB2* expression. We defined a transcriptional signature discriminating the two CD8+ populations, including differential expression of key residency-associated genes and cytotoxic molecules. Flow cytometry of recipient-derived cells infiltrating the graft and intestinal lymphocytes from healthy gut confirmed the two CD8+ T_RM_ phenotypes, with β2-integrin acting as a CD103-CD8+ T_RM_ marker. CD103+ CD8+ T_RM_ cells produced IL-2, and demonstrated greater polyfunctional cytokine production, while β2-integrin+ CD69+ CD103-T_RM_ cells had higher granzyme expression. Phenotypic and functional analysis of intestinal CD4+ T cells identified many parallels, including a distinct β2-integrin+ population. Together, these results describe the transcriptional, phenotypic, and functional heterogeneity of human intestinal T_RM_ cells, and suggest a role for β2-integrin in T_RM_ development.

**Summary:** Heterogeneity within human tissue-resident memory T (T_RM_) cells is poorly understood. We show that transcriptionally, phenotypically, and functionally distinct CD4+ and CD8+ T_RM_ subsets exist in the human intestine, and that β2-integrin expression identifies a distinct population of CD8+ T_RM_ cells.

## Introduction

Tissue-resident memory T (T_RM_) cells are a subset of long-lived T cells that reside in tissue and do not recirculate (Mackay and Kallies, 2017; Szabo et al., 2019). T_RM_ populations provide rapid, *in situ* adaptive protection against a wide spectrum of pathogens (Gebhardt et al., 2011; Schenkel et al., 2014). T_RM_ cells also have key roles in cancer immune surveillance (Park et al., 2019), and are implicated in autoimmunity, including inflammatory bowel disease (IBD) and coeliac disease (Zundler et al., 2019; Mayassi et al., 2019). CD8+ T_RM_ cells have potent cytotoxic functions and produce pro-inflammatory cytokines to trigger innate and adaptive immune responses (Ariotti et al., 2014).

Murine work has advanced our understanding of T_RM_ cells substantially, however T_RM_ phenotype, transcriptional profiles, and genetic regulation differ between mice and humans (Oja et al., 2017; Hombrink et al., 2016; Kumar et al., 2017). Until recently, human studies of T_RM_ biology were hampered by the inability to prove long-lived tissue residency, with surface molecules CD69 and CD103 (αE integrin) used as surrogate T_RM_ markers. These were used to identify the putative transcriptional signature of human T_RM_ cells, with CD69 hypothesised as the key residency marker (Kumar et al., 2017). However, this gene signature was derived from bulk populations, so it remains unclear if there are transcriptionally distinct subsets within human T_RM_ cells.

Despite its expression on almost all murine and human T_RM_ cells, the use of CD69 as a residency marker has recently been questioned. CD69 restricts lymphocyte tissue egress via S1P1 inhibition, but is not required for the development of functional T_RM_ cells in mice (Shiow et al., 2006; Walsh et al., 2019). CD69 can be induced via stimulation, and a proportion of CD69+ T cells in tissue are not resident, making CD69 a suboptimal residency marker (Shiow et al., 2006; Sancho et al., 2005; Beura et al., 2018). Therefore, additional phenotypic markers to identify CD103-T_RM_ populations are required.

Recent work has exploited the human model of organ transplantation to study long-lived donor-derived T cells, which are definitive functionally resident T_RM_ cells (Zuber et al., 2016; Bartolomé-Casado et al., 2019; Snyder et al., 2019). Studies using this approach have demonstrated persistence of clonally-identical intestinal CD8+ T_RM_ cells for up to 1 year in the small intestine (SI) (Bartolomé-Casado et al., 2019), and of donor-derived T cells for 600 days post-transplant (Zuber et al., 2016). This approach was also used to identify putative SI CD8+ T_RM_ cell subsets based on expression of CD103 and KLRG1, with differences in clonality, granzyme expression, and cytokine production (Bartolomé-Casado et al., 2019). However, it remains unclear whether these cell populations represent transcriptionally distinct subsets.

This work sought to examine the heterogeneity within functionally resident donor-derived T cells in intestinal transplantation using flow cytometry and single-cell RNA sequencing (scRNAseq). We confirmed that SI CD4+ and CD8+ T_RM_ cells can persist for 5 years post-transplant. scRNAseq identified conventional and regulatory CD4+ T_RM_ cell populations, as well as two transcriptionally distinct CD8+ T_RM_ subsets, which differed in expression of *ITGAE* (CD103, αE integrin) and *ITGB2* (CD18, β2-integrin). These two populations differentially expressed putative T_RM_-associated genes, indicating that the gene signatures derived from bulk RNAseq data may be a synthesis of several transcriptomic profiles. We validated this phenotypic and functional heterogeneity in the healthy intestine, with increased β2-integrin expression and distinctive effector function in CD103-T_RM_ cells. CD69, β2-integrin, and CD103 expression altered with time post-transplant on recipient-derived graft-infiltrating CD8+ T cell populations, consistent with acquisition of T_RM_ status. We conclude that CD69+ CD103-β2-integrin+ CD8+ intestinal T cells are a transcriptionally and functionally distinct T_RM_ population, and suggest that β2-integrin can serve as a proxy surface marker for the CD103-CD8+ T_RM_ population.

## Results and discussion

### Long-lived conventional CD4+ and CD8+ T cells, but not innate-like populations, persist long-term in the human intestine

To examine the persistence of resident T cells in the small intestine (SI) post-transplant, we used HLA allele congenic cell tracking, a method allowing discrimination of donor- and recipient-derived cells following transplantation using fluorophore-conjugated antibodies to discordant Class I HLA haplotypes (Fig. 1A, Supplemental Fig. 1A) (Zuber et al., 2016; Bartolomé-Casado et al., 2019). The presence of SI donor- and recipient-derived T cells *in situ* was confirmed by chip cytometry (Fig. 1B)(Leng et al., 2019).

**Figure 1.**
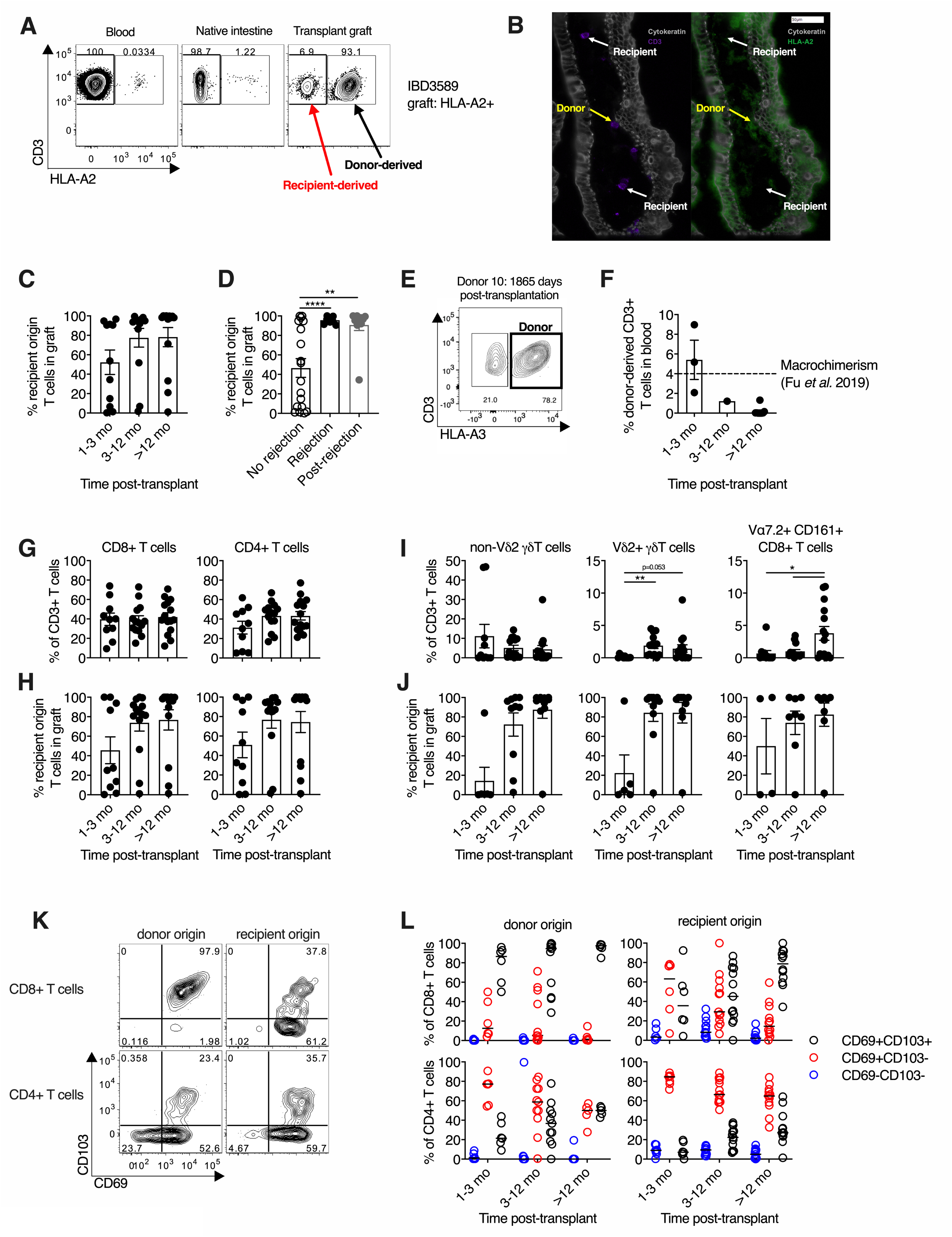
Conventional T cell subsets persist for 5 years post-transplant, and display a characteristic tissue resident memory (T_RM_) phenotype. (A) Representative flow cytometry plot of HLA-A2 expression of T cells from the blood, the recipient native intestinal mucosa, and the intestinal transplant graft, demonstrating identification of donor- and recipient-derived populations by HLA mismatch. (B) Representative chip cytometry image of intestinal graft mucosa demonstrating presence of donor-derived (HLA-A2+, yellow arrows) and recipient-derived (HLA-A2-, white arrows) CD3+ T cells in the lamina propria. Cytokeratin (grey); CD3 (purple); HLA-A2 (green). (C) Percentage of recipient-origin CD3+ T cells in intestinal grafts, categorised by time post-transplant (n=37; mean +/- SEM). (D) Percentage of recipient-origin CD3+ T cells in intestinal grafts, categorised by history of graft rejection. Unfilled circles, no rejection; Black, current rejection; Grey, previous (post) rejection. (n=37; mean +/- SEM). (E) Flow cytometry plot of HLA-A3 expression of graft-derived T cells in one subject who demonstrated persistent donor chimerism in the intestinal graft 1865 days (5 years and 1 month) post-transplant. (F) Percentage of donor-origin CD3+ T cells in the blood of intestinal transplant recipients, categorised by time post-transplant (n=14; mean +/- SEM). Dashed line at 4% represents the cut-off for significant macrochimerism from prior studies (Fu et al., 2019; Zuber et al., 2015). (G) Conventional CD8+ and CD4+ T cell subsets in the small intestinal graft as a proportion of total T cells, categorised by time post-transplant (n=41; mean +/- SEM). (H) The percentage of recipient-derived T cells infiltrating the intestinal graft within conventional CD8+ and CD4+ T cell subsets, categorised by time post-transplant (n=41; mean +/- SEM). (I) Unconventional non-Vδ2+ γδ T cell, Vδ2+ γδ T cell, and Vα7.2+CD161+ CD8+ T cell (mucosal-associated invariant T cells) subsets in the small intestinal graft as a proportion of total T cells, categorised by time post-transplant (n=41; mean +/- SEM). (J) The percentage of recipient-derived T cells infiltrating the intestinal graft within unconventional non-Vδ2+ γδ T cell (n=27), Vδ2+ γδ T cell (n=25), and Vα7.2+CD161+ CD8+ T cell subsets (20), categorised by time post-transplant (mean +/- SEM). (K) Representative flow cytometry plot of CD69 and CD103 expression on donor- and recipient-derived CD8+ and CD4+ T cells in the intestinal graft. (L) Percentage of donor- and recipient-derived CD8+ and CD4+ T cells in the intestinal graft co-expressing CD69 and CD103 categorised by time post-transplant (n=35; median marked with black line). Blue, CD69-103-cells; Red, CD69+CD103-cells; Black, CD69+CD103+ cells. For further analysis of surface marker expression, rare populations with fewer than 10 cells were excluded from the analysis (J, L). Statistical analysis performed with 1-way ANOVA with Dunnett’s multiple comparisons test. *, P ≤ 0.05; **, P ≤ 0.01.

Infiltration of recipient-derived T cells into the graft increased over time, with striking heterogeneity (Fig. 1C). Current or previous rejection was associated with higher proportions of recipient-derived T cells in the graft (Fig. 1D), consistent with prior work (Zuber et al., 2016). Conversely, two individuals had persistent donor-derived SI T cell populations at 4 years 7 months (1684 days) and 5 years 1 month (1865 days) post-transplant (Fig. 1E, and data not shown). This extends previous reports of long-lived donor-chimerism in the human intestine (Zuber et al., 2016; Bartolomé-Casado et al., 2019).

Haematopoietic stem and progenitor cells can persist in the intestinal transplant graft, sometimes leading to long-term chimerism of donor-derived populations in blood (Fu et al., 2019). This could allow persistence of donor-derived T cells in the graft through continuous ingress from blood, rather than via residency (Bartolomé-Casado et al., 2019). Contrasting with the work of Fu *et al*., flow cytometric analysis of blood collected from transplant recipients at the time of biopsy revealed the percentage of circulating donor origin T cells was extremely low beyond 3 months post-transplant (median 0.042%, 95% CI 0-1.22%; Fig. 1F). This is far below the posited cut-off for macrochimerism of 4% in previous work (Fu et al., 2019), and similar to background non-specific staining in non-transplant recipients (0.01-0.12%; Supplemental Fig. 1B), suggesting against continuous replacement as a confounding mechanism for sustained donor chimerism. This discrepancy in circulating chimerism between studies may be due to differences in transplant procedure (multi-visceral vs isolated intestinal transplant in this study) and recipient age (paediatric vs adult in this study) (Zuber et al., 2015; Fu et al., 2019).

Conventional CD4+ and CD8+ T cells dominated in the graft post-transplant (Fig. 1G), and demonstrated similar kinetics of recipient-derived replacement with time (Fig. 1H). We also examined the dynamics of unconventional T cell subsets post-transplant, as their residency characteristics in humans are poorly understood (Supplemental Fig. 1C). CD161+Vα7.2+CD8+ T cells (consistent with mucosal-associated invariant T (MAIT) cells), and Vδ2+ γδ T cells (which possess analogous innate-like functions to MAIT cells (Provine et al., 2018; Gutierrez-Arcelus et al., 2019)) were rare in the graft post-transplant, recovering to expected frequencies after 1 year (Fig, 1I). However, the non-Vδ2 γδ T cell subsets demonstrated different dynamics, with this population present at early timepoints. Unconventional T cell subsets demonstrated similar replacement kinetics (Fig. 1J). It is unclear if the low innate-like T cell frequency post-transplant is due to differences in residency characteristics, or to increased sensitivity to the ischaemic insult of surgery or perioperative conditioning regimes.

We examined CD69 and CD103 expression on donor- and recipient-derived CD4+ and CD8+ T cell populations in the intestinal graft. CD103 expression was restricted to CD69+ cells, with a greater proportion of CD8+ cells expressing CD103 than CD4+ cells(Fig. 1K), in keeping with prior work (Kumar et al., 2017). Donor-derived T cells showed near-ubiquitous expression of CD69 consistent with a lack of recent migration from blood, and had fewer CD69-cells than recipient-derived populations for both CD8+ (median 0.03% vs 4.17%, p<0.0001, Wilcoxon signed rank test) and CD4+ cells (median 0.10% vs 5.65%, p<0.0001, Wilcoxon signed rank test) (Fig. 1L). This suggests that CD69-T cells in the SI are not functionally resident in humans, in contrast to recent murine data which showed no functional requirement for CD69 to establish intestinal residency (Walsh et al., 2019). Recipient-derived CD4+ and CD8+ T cell populations showed increasing expression of CD103 with time, consistent with the acquisition of a T_RM_ phenotype, as in prior work (Zuber et al., 2016). We have previously shown higher expression of the C-type lectin-like receptor CD161 on intestinal CD103+ CD8+ T cells (Fergusson et al., 2015); here a greater proportion of donor-derived CD8+ T cells expressed CD161, consistent with an association with residency (Supplemental Fig. 1D,E).

### Single-cell RNA sequencing delineates transcriptionally distinct states within CD4+ and CD8+ T_RM_ populations

The persistence of CD103- and CD103+ donor-derived T cells up to five years post-transplant, and the enrichment of CD103+ recipient-derived T cells at later time points, raised the possibility that CD103- and CD103+ T cells represented distinct cell states. To test this hypothesis, we performed droplet-based scRNAseq of sorted donor-derived graft-resident T_RM_ cells from a single subject one year post-transplant (Experiment 1, Fig. 2A). 1,774 cells were captured and sequenced, with 974 cells remaining after filtering (Supplemental Fig. 2A).

**Figure 2.**
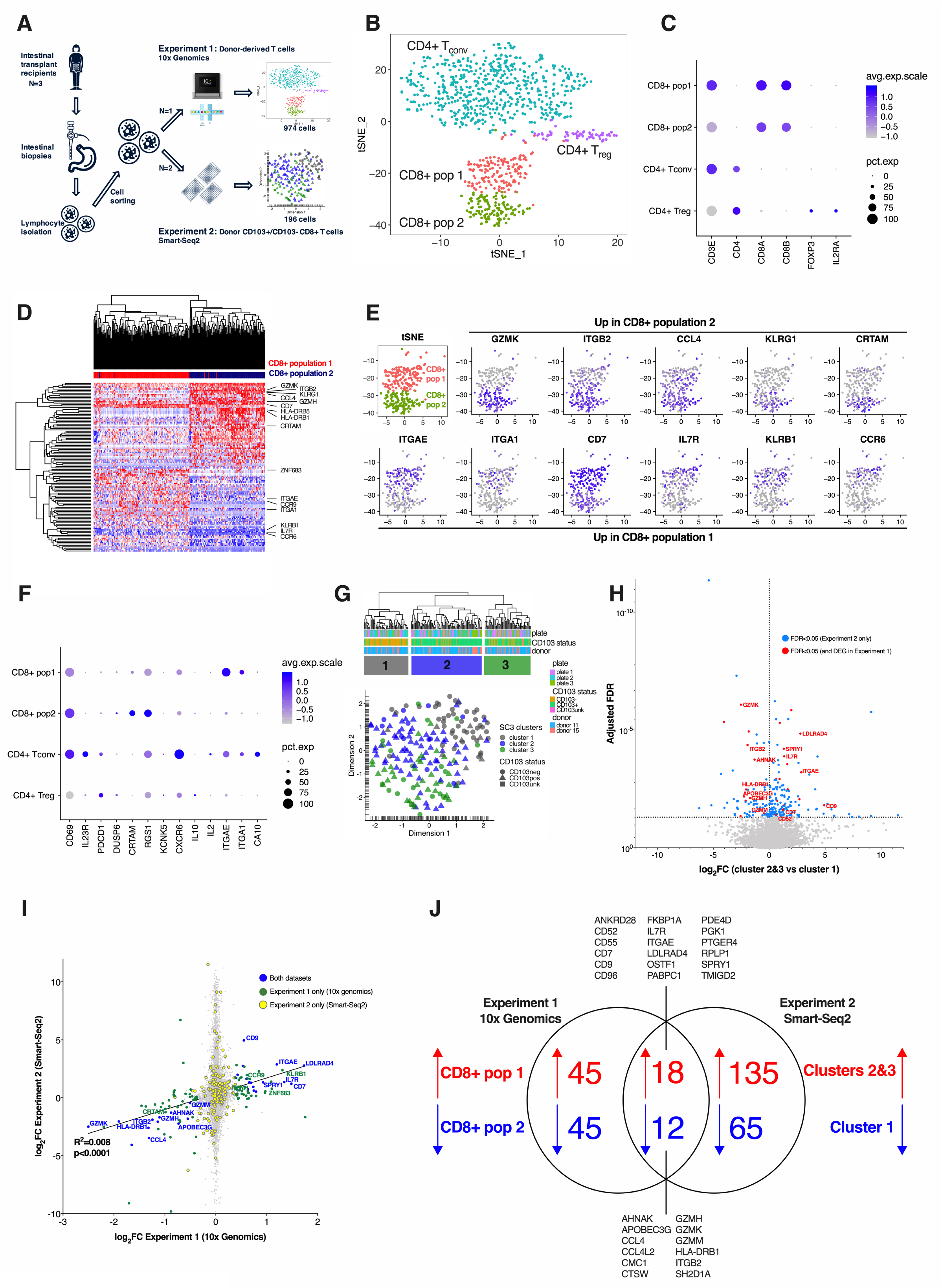
Transcriptionally distinct tissue resident memory (T_RM_) T cell populations at single cell level, delineated by CD103 expression. (A) Schematic diagram of two scRNAseq experiments. Biopsies of small intestinal transplant tissue were collected from subjects at endoscopy, then dissociated to isolate intestinal lymphocytes. These were sorted by FACS, first with bulk sorting of donor-derived T cells in Experiment 1, then with index sorting of donor-derived CD103- and CD103+ T cells in Experiment 2, before scRNAseq library preparation and sequencing using the 10x Genomics platform (Experiment 1) or the Smart-Seq2 protocol (Experiment 2). (B) tSNE plot of 974 donor-derived T cells from a single subject showing four transcriptionally distinct populations of conventional CD4+ and CD8+ T cells. (C) Dot plot of key gene identifiers for the four clusters showing two clusters of CD8+ T cells, and two clusters of CD4+ T cells, including one containing cells expressing *IL2RA* and *FOXP3*, consistent with a regulatory T cell phenotype. Dot size indicates the proportion of cells in which the gene is present. Colour intensity indicates the mean expression level of the gene in cells in that cluster. (D) Heatmap indicating hierarchical clustering of gene expression of CD8+ donor-derived T cells from population 1 and population 2. Cell labels above indicate cluster: Red, population 1; Blue, population 2. Genes of interest are highlighted. (E) tSNE plot showing the expression of key genes upregulated in CD8+ population 2 (top row) or upregulated in CD8+ population 1 (bottom row). (F) Dot plot showing the expression of 13 genes previously associated with tissue residency in human CD69+ T cells (Kumar et al., 2017). Dot size indicates the proportion of cells in which the gene is present. Colour intensity indicates the mean level of expression of the gene in cells in that cluster. (G) Hierarchical clustering and UMAP plot of 196 index-sorted CD103- and CD103+ CD8+ donor-derived T cells from two subjects identified three transcriptionally distinct clusters of conventional CD8+ T cells. Cell labels below the dendrogram indicate the plate used for sorting, the subject, and the CD103 expression determined by index sorting. For UMAP plot: circles, CD103-cells by index-sorting; triangles, CD103+ cells by index-sorting; square, unknown CD103 expression by index-sorting. (H) Volcano plot of differential gene expression between cluster 1 and clusters 2 & 3 identified in (G), with genes upregulated (right) or down regulated (left) in clusters 2&3 compared to cluster 1. log_2_(fold change) is plotted against the FDR-adjusted p-value, with horizontal dotted line at FDR=0.05. Differentially expressed genes are marked in blue, with those differentially expressed both Experiments 1 and 2 marked in red. (I) Correlation of log_2_(fold change) between populations 1 vs 2 from Experiment 1 (10x Genomics, x-axis) and clusters 2&3 vs cluster 1 from Experiment 2 (Smart-Seq2, y-axis). Blue, genes differentially expressed in both experiments; green, genes differentially expressed in Experiment 1 only; yellow, genes differentially expressed in Experiment 2 only; grey, genes not differentially expressed. Genes shown were expressed in both experiments. (J) Venn diagram showing differentially expressed genes upregulated (red) or downregulated (blue) in cluster 1 (CD103-) of Experiment 2 (Smart-Eeq2), or CD8+ population 2 of Experiment 1 (10x Genomics). A core gene set of 30 genes that differentiated the two populations in both experiments are listed.

Four transcriptionally distinct clusters were identified (Fig. 2B): conventional CD4+ T cells, regulatory CD4+ T cells expressing *IL2RA* and *FOXP3*, and two clusters of conventional CD8+ T cells (Fig. 2C). The differentially expressed genes (DEGs) between the two populations of donor-derived CD8+ T_RM_ cells (hereafter CD8+ population 1 and 2) were analysed (Fig. 2D). Population 1 expressed *ITGAE* (CD103), as well as higher levels of *CD7*, *IL7R* (CD127), *KLRB1* (CD161), and the chemokine receptor *CCR6* (Fig. 2E). IL-7, a stromal-derived homeostatic cytokine which provides survival and proliferative signals to lymphocytes (Raeber et al., 2018), is required for epidermal T_RM_ persistence (Adachi et al., 2019), and *CD127* is highly expressed in SI memory T cells (Thome et al., 2014). The differential expression of IL7R suggests differences in the mechanisms, or nature, of T_RM_ persistence between the two populations (Raeber et al., 2018).

Conversely, CD8+ population 2 expressed low levels of *ITGAE*, but high levels of *KLRG1*, as well as cytotoxic granzyme molecules and class II MHC molecules (Fig. 2D,E). Surface KLRG1 expression can delineate two putative SI CD103-CD8+ T cell subsets with similar residency characteristics (Bartolomé-Casado et al., 2019). However, population 2 did not sub-cluster further based on *KLRG1* gene expression, suggesting that these are not transcriptionally distinct states. Of particular interest, the integrin *ITGB2* (β2-integrin/CD18), was highly expressed by the CD103-CD8+ population 2. β2-integrin can form heterodimers with four α-integrins (Fagerholm et al., 2019), only one of which, *ITGAL* (CD11a), was detected in the dataset. *ITGAL* was highly expressed on the CD103-CD8+ population 2 (Supplemental Fig. 2B,C).

### CD8+ CD103+ and CD103-ITGB2hi T_RM_ subsets differ in expression of putative residency-associated genes

Transcriptional signatures associated with human tissue residency have been defined, most thoroughly by bulk RNA sequencing of CD69- and CD69+ T cells from multiple tissues (Kumar et al., 2017). This T_RM_ gene set was explored in the two CD8+ T cell populations. The genes downregulated in CD69+ T cells were either not detected, or found at low levels in both clusters (Supplemental Fig. 2D,E). However, several T_RM_-associated genes upregulated in CD69+ T cells were differentially expressed between the two clusters, with population 1 expressing higher levels of *ITGAE* and *ITGA1* (in agreement with work on renal T_RM_ cells (de Leur et al., 2019)), and population 2 expressing higher levels of *CRTAM* (Fig. 2F, Supplemental Fig. 2F). In addition, *ZNF683* (Hobit), a transcriptional regulator of residency in mice (Mackay et al., 2016), was more highly expressed in population 1. These data indicate that previously identified T_RM_ gene signatures may represent an amalgamation of several distinct T_RM_ transcriptional states.

To confirm the presence of transcriptionally distinct CD8+ T_RM_ states, a second scRNAseq experiment was performed on samples from two further transplant recipients (Experiment 2, Fig. 2A). Donor-derived CD103- and CD103+ CD8+ T cells were index-sorted before plate-based scRNAseq using the Smart-Seq2 protocol (Picelli et al., 2013). 267 cells were sorted and sequenced, with 196 cells remaining post-filtering (Supplemental Fig. 2G-I). Three clusters were identified, with cluster 1 predominantly formed of CD103-T cells, and the transcriptionally similar clusters 2 and 3 formed of CD103+ T cells (Fig. 2G). Clusters 2 and 3 (CD103+) expressed higher levels of *ITGAE*, *CD7*, and *IL7R*, while cluster 1 (CD103-) expressed higher levels of *GZMK*, *GZMH*, class II HLA molecules, and *ITGB2* (Fig. 2H). *ITGAL*, was also detected in cluster 1 (Supplemental Fig. 2J).

Differential expression analysis between cluster 1 and clusters 2 and 3 combined revealed similar transcriptional differences to those in Experiment 1 (Fig. 2I, Supplemental Table 2). Comparison of DEGs in the two experiments identified a core set of 30 genes that distinguished the two populations (Fig. 2I,J).

Putative T_RM_ cell subsets in the context of lung transplantation have been described, however these did not align with CD103 expression (Snyder et al., 2019), in contrast to our work. The transcriptional signatures of the clusters were also different in our work, with the exception of *ZNF683*, which was also associated with one lung T_RM_ subset. These differences may reflect tissue-specific gene signatures in T_RM_ cells, differences in CD4+ and CD8+ T_RM_ cells, which co-clustered in the previous study of lung T_RM_ cells, or may be an effect of increased cell number in this study, allowing greater power to detect transcriptionally distinct sub-clusters.

### CD103+ and CD103-CD8+ T cells display distinct phenotypes in the healthy intestine

The presence of CD103- and CD103+ donor-derived CD8+ T cells in the transplanted SI mucosa was confirmed using chip cytometry (Fig. 3A). Flow cytometry of donor-derived T cells demonstrated differences in expression of CD161, β2 integrin, and granzyme K between CD103- and CD103+ CD8+ T cells, consistent with scRNAseq data (Supplementary Fig. 3A).

**Figure 3.**
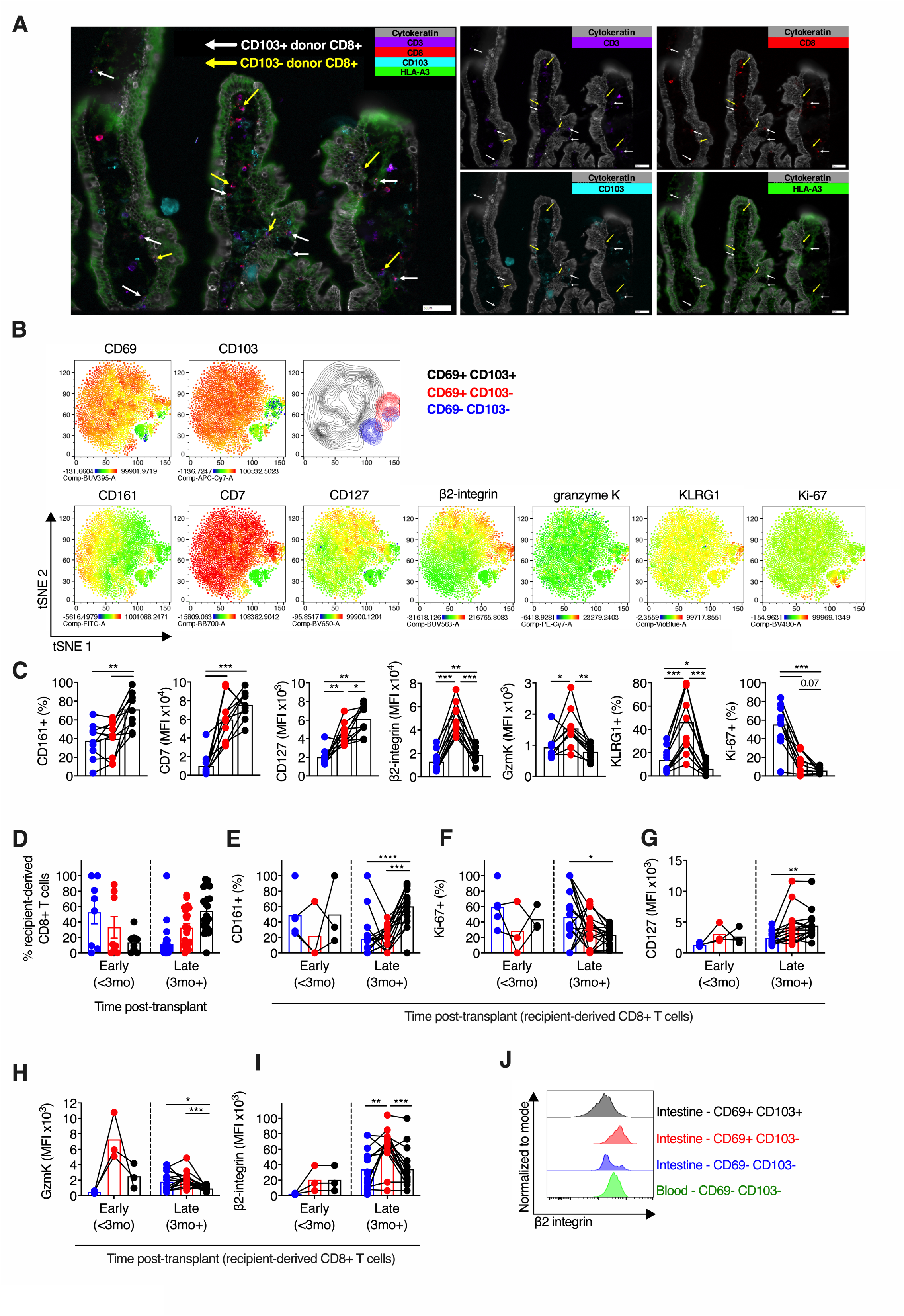
Localisation and phenotype of CD103- and CD103+ intestinal T_RM_ CD8+ populations. (A) Fluorescence microscopy chip cytometry image of small intestinal transplant mucosa from a single subject 3 months post-transplant. The donor was HLA-A3+, and recipient was HLA-A3-. False colour fluorescence imaging for cytokeratin (grey), CD3 (purple), CD8 (red), CD103 (blue), and HLA-A3 (green). Donor-derived CD8+ CD103+ and CD103-cells are indicated by white and yellow arrows respectively. (B,C) Phenotypic analysis of CD8+ T cell populations in healthy small intestine. (B) tSNE representation of CD8+ T cell phenotype from spectral flow cytometry concatenated from 5 healthy control subjects, with expression of the following markers superimposed: CD69 and CD103 (first row); CD161, CD7, CD127, β2-integrin, granzyme K, KLRG1, and Ki-67 (second row). The location of cells co-expressing CD69 and CD103 is indicated. Blue, CD69-103-cells; Red, CD69+CD103-cells; Black, CD69+CD103+ cells. (C) Proportion of positive cells (CD161, KLRG1, Ki-67) or MFI (CD7, CD127, β2-integrin, granzyme K) of CD8+ T cells, categorised by CD69 and CD103 expression, in small intestinal biopsies from healthy control subjects (n=10). Mean percentage or MFI represented by bars. Connecting lines represent populations from the same subject. Blue, CD69-103-cells; Red, CD69+CD103-cells; Black, CD69+CD103+ cells. (D-J) Phenotypic analysis of recipient-derived CD8+ T cell populations infiltrating the intestinal transplant graft. (D) The proportion of recipient-derived CD8+ T cells co-expressing CD69 and CD103 in intestinal transplant grafts, categorised by time post-transplant (n=35). Blue, CD69-103-cells; Red, CD69+CD103-cells; Black, CD69+CD103+ cells. (E-I) Proportion of CD161+ cells (E), Ki-67+ cells (F) or MFI of CD127 (G), granzyme K (H), and β2 integrin (I) of recipient-derived CD8+ T cells, categorised by CD69 and CD103 expression and time post-transplant, in intestinal transplant grafts (n=23). Mean percentage or MFI represented by bars. Black lines connect populations from the same sample. Blue, CD69-103-cells; Red, CD69+CD103-cells; Black, CD69+CD103+ cells. (J) Representative histogram of β2 integrin surface expression by circulating, intestinal CD69-, intestinal CD69+CD103-, and intestinal CD69+CD103+ CD8+ T cells. Statistical analysis performed with one-way ANOVA with Tukey’s multiple comparisons test. *, P ≤ 0.05; **, P ≤ 0.01; ***, P ≤ 0.001; ****, P ≤ 0.0001.

To validate the phenotypic differences between the two putative SI CD8+ T_RM_ subsets outside the transplant setting, we performed flow cytometry of SI T cells from healthy donors. CD8+ SI T cells expressing both CD69 and CD103 predominated, representing 88.9% (81.7-96.0%) of CD8+ T cells, with no difference seen between SI location (Supplementary Fig. 3B).

CD103+ CD8+ T cells expressed higher levels of CD161 and CD127 (IL7R) compared with the CD69- or CD69+CD103-cells, consistent with the transcriptomic data (Fig. 3B,C). CD7 expression was higher on all CD69+ T cells, with no difference seen between CD69+CD103- and CD69+CD103+ populations. In contrast, CD69+CD103-CD8+ T cells expressed higher levels of β2-integrin, granzyme K, and KLRG1 than either CD69-cells or CD69+CD103+ cells.

Ki-67 expression formed a gradient between the three populations, with higher expression in the CD69-population, and a trend towards increased Ki67 expression in CD69+CD103-CD8+ T cells compared to CD103+ CD8+ T cells (mean 14.88% vs 5.72%, 1-way ANOVA and Tukey’s multiple comparisons test, P=0.067). This is consistent with prior work indicating that T_RM_ cell persistence is due to longevity rather than *in situ* proliferation (Thome et al., 2014).

### Graft-infiltrating, recipient-derived T cells take on a T_RM_ phenotype over time, with CD103+ and CD103-CD8+ T cells displaying distinct phenotypes

To explore the dynamics of T_RM_ phenotype acquisition, we examined graft-infiltrating, recipient-derived T cells. In the early post-transplant period, 52.7% of infiltrating CD8+ T cells lacked CD69 expression, with acquisition of the CD69+ CD103+ T_RM_ phenotype over time (Fig. 3D). At later times post-transplant, CD103- and CD103+ populations clearly differed in phenotype, consistent with the two subsets seen in healthy SI. CD161 expression was higher on CD103+ CD8+ T cells than CD69+CD103-T cells, and granzyme K expression was higher on CD69+CD103-CD8+ T cells (Fig. 3E-H). A similar gradient of Ki-67 expression between the populations was seen (Fig. 3F). In addition, graft-infiltrating CD69+CD103-CD8+ T cells demonstrated higher expression of β2 integrin than either CD69- or CD103+ populations (Fig. 3I). β2 integrin expression is constitutively high on circulating T cells (Fig. 3J), as LFA-1 is involved in tissue entry via ICAM-1 (Fagerholm et al., 2019). These data suggest that β2 integrin surface expression is reduced on recent tissue immigrants, before subsequent up regulation on CD69+CD103-T_RM_ cells, but not on CD103+ T_RM_ cells. Recipient-derived CD8+ T cells infiltrating the graft in the early post-transplant period displayed less clear distinctions between populations, although the small number of samples precluded further analysis.

### CD103+ CD8+ intestinal T cells demonstrate greater capacity for cytokine production

To assess cytokine production capacity of these subsets, SI T cells from healthy controls were stimulated for four hours with PMA+ionomycin, before intracellular flow cytometry for TNFα, IFNγ, IL-2, CCL4, IL-17A, and IL-10. 77% of CD103+ CD8+ T cells produced at least one cytokine, a greater proportion than CD69+CD103-CD8+ T cells (47.7% cytokine positive, P<0.0001), and CD69-CD8+ T cells (37% cytokine positive, P<0.0001) (Fig. 4A-F). CD69+ populations produced more TNFα and IFNγ than CD69-cells, irrespective of CD103 expression (Fig. 4A,B). While scRNAseq data indicated increased *CCL4* transcripts in CD69+CD103-CD8+ T cells, CCL4 production following stimulation was not different between the two CD69+ populations (Fig. 4C).

**Figure 4.**
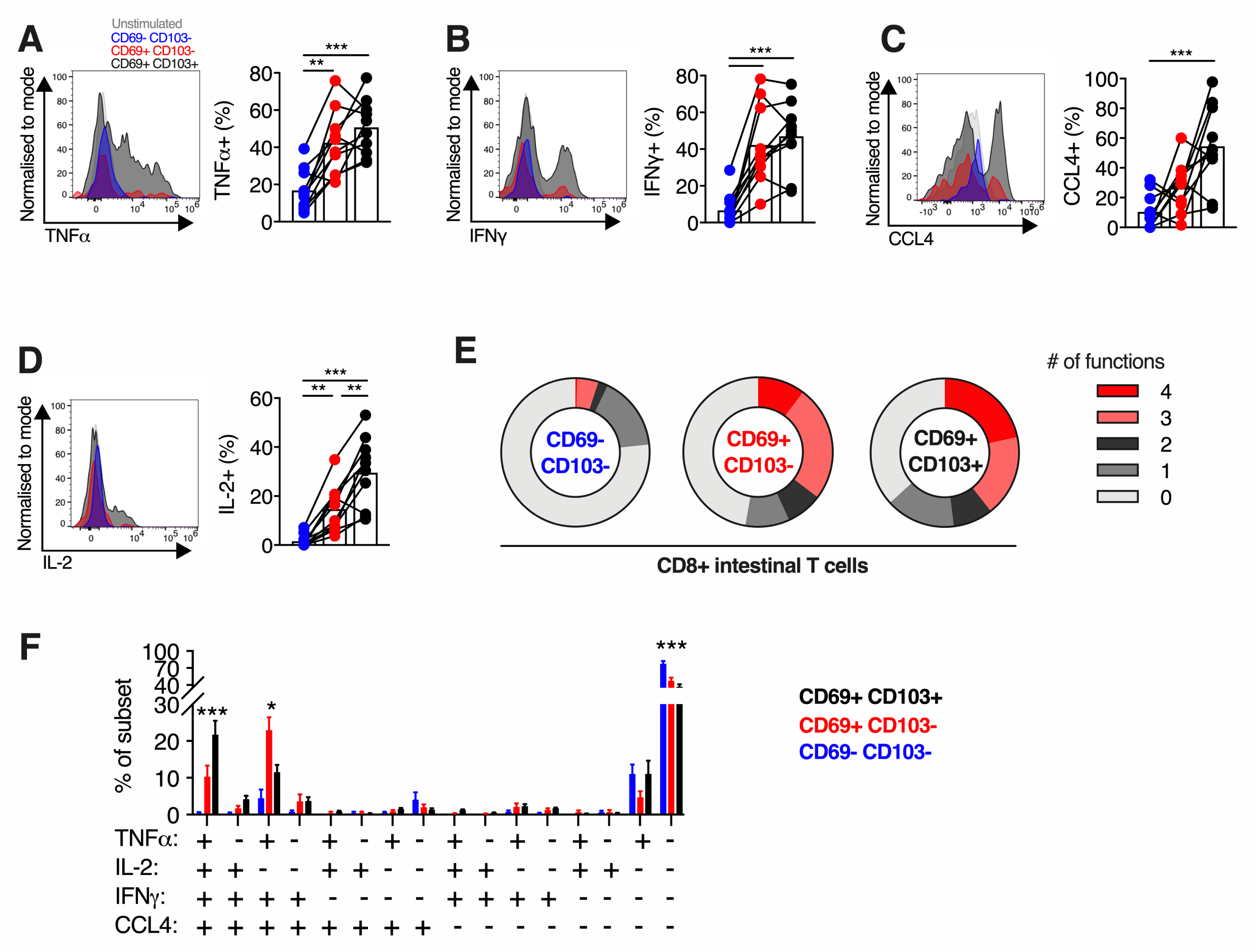
Functional capacity of small intestinal CD8+ T cell populations. (A-D) Cytokine production by small intestinal CD8+ T cells. Representative histograms of expression, and group summaries of proportion of CD8+ T cells expressing TNFα (A), IFNγ (B), CCL4 (C), and IL-2 (D) after 4-hour stimulation with PMA and ionomycin in the presence of brefeldin A and monensin, categorised by CD69 and CD103 expression, in small intestinal biopsies from healthy control subjects (n=10). Blue, CD69-103-cells; Red, CD69+CD103-cells; Black, CD69+CD103+ cells. (E) Mean proportion of CD8+ T cells expressing 0, 1, 2, 3, or 4 of the cytokines or chemokines TNFα, IFNγ, CCL4, or IL-2, categorised by CD69 and CD103 co-expression, from small intestinal biopsies from healthy control subjects (n=10). (F) Mean percentage (+/- SEM) of CD8+ T cells co-expressing TNFα, IFNγ, CCL4, or IL-2 production after PMA/ionomycin stimulation as described, categorised by CD69 and CD103 expression. Blue, CD69-103-cells; Red, CD69+CD103-cells; Black, CD69+CD103+ cells. Statistical analysis performed with one-way ANOVA with Tukey’s multiple comparisons test. *, P ≤ 0.05; **, P ≤ 0.01; ***, P ≤ 0.001.

CD103+ CD8+ T cells expressed more IL-2 than either CD69+CD103- or CD69-populations (Fig. 4D). Hepatic CD103+ CD8+ T_RM_ cells also produce increased IL-2 (Pallett et al., 2017). Autocrine IL-2 production is critical to secondary proliferative responses and IFNγ production in CD8+ T cells (Feau et al., 2011), and may be of particular relevance to intra-epithelial lymphocyte populations, where the infrequent CD4+ T cells may provide suboptimal help (Zimmerli et al., 2005).

There was a spectrum of functionality between the three populations, with CD69-CD8+ T cells predominantly non- or mono-functional, CD103+ CD8+ T cells predominantly polyfunctional, and CD69+CD103-CD8+ T cells showing intermediate functionality (Fig. 4E,F), consistent with a prior report (Bartolomé-Casado et al., 2019). Quadruple functional cells were more common in the CD103+ population than in the CD69+CD103-population (21.76% vs 10.32%, P<0.0001), and were near-absent in the CD69-population (0.38%). These results demonstrate that the transcriptionally distinct CD103- and CD103+ CD8+ T_RM_ populations differ functionally and phenotypically, with CD103+ CD8+ populations more polyfunctional, and producing IL-2.

### CD103+ and CD103-CD4+ T cells display analogous phenotypic and functional differences to their CD8+ counterparts

Despite not forming transcriptionally distinct clusters by scRNAseq, CD4+ SI T cells differed in their phenotype, dependent on CD69 and CD103 expression. CD69-populations expressed lower levels of CD161 and CD127, while CD69+CD103-CD4+ T cells had higher expression of β2-integrin than either CD69- or CD103+ populations, analogous to their CD8+ counterparts (Fig. 5A). Recipient-derived, graft-infiltrating CD69+CD103-CD4+ T cells also displayed higher expression of β2-integrin than CD103+ counterparts (MFI 51,753 vs 30,959, P<0.0001; Fig. 5B).

**Figure 5.**
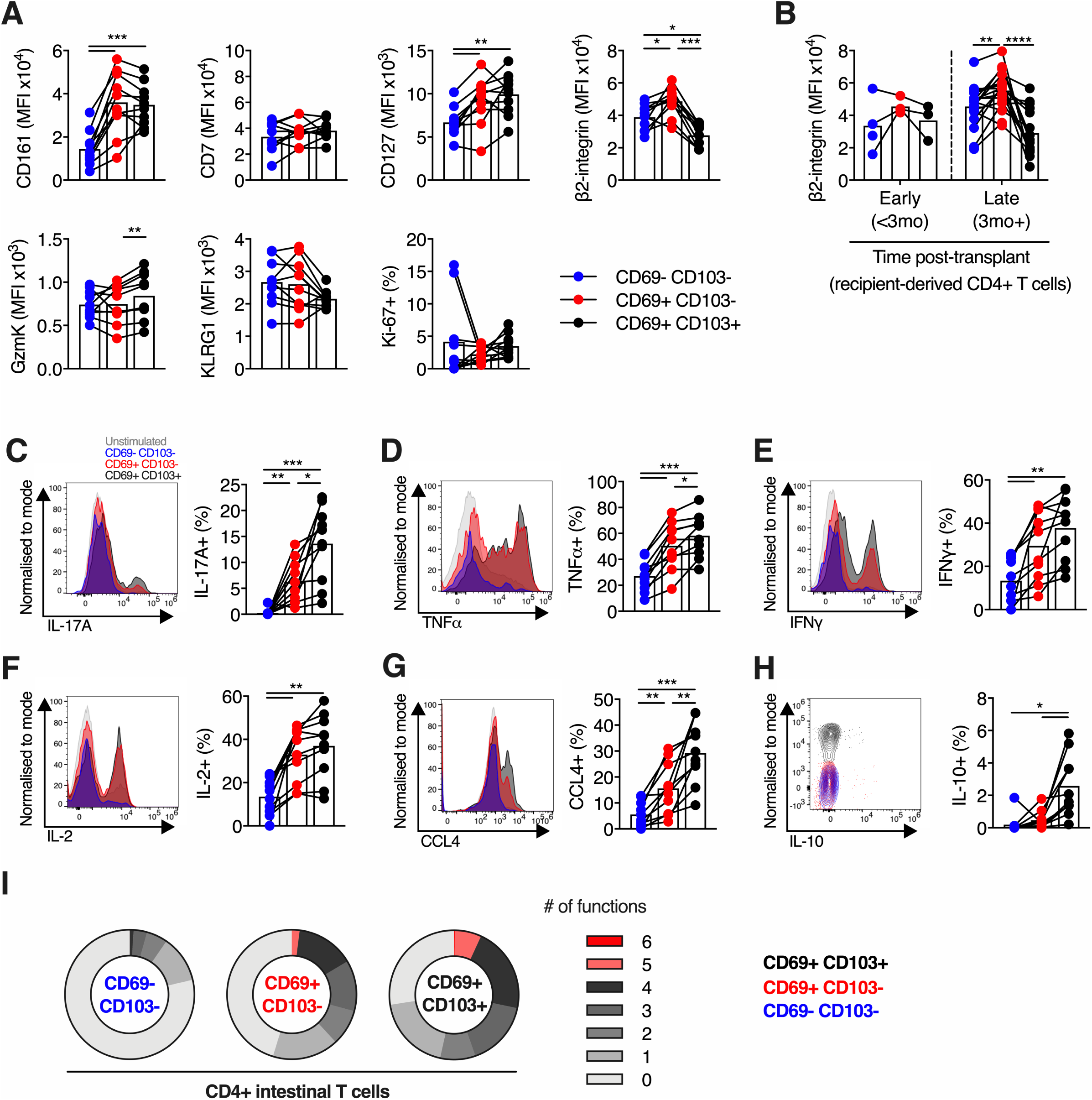
Phenotype and function of small intestinal CD4+ T cell populations. (A) MFI (CD161, CD7, CD127, β2-integrin, granzyme K, and KLRG1) or percentage positive (Ki67) of CD4+ T cells, categorised by CD69 and CD103 expression, in small intestinal biopsies from healthy control subjects (n=10). MFI represented by bars. Connecting lines represent populations from the same subject. Blue, CD69-103-cells; Red, CD69+CD103-cells; Black, CD69+CD103+ cells. (B) MFI of β2-integrin of recipient-derived CD4+ T cells, categorised by CD69 and CD103 expression and time post-transplant, in intestinal transplant grafts (n=21). MFI represented by bars. Blue, CD69-103-cells; Red, CD69+CD103-cells; Black, CD69+CD103+ cells. (C-I) Cytokine production by small intestinal CD4+ T cells. Representative histograms of expression, and group summaries of proportion of CD4+ T cells expressing IL-17A (C), TNFα (D), IFNγ (E), IL-2 (F), CCL-4 (G), and IL-10 (H) after 4-hour stimulation with PMA and ionomycin in the presence of brefeldin A and monensin, categorised by CD69 and CD103 expression, in small intestinal biopsies from healthy control subjects (n=10). Blue, CD69-103-cells; Red, CD69+CD103-cells; Black, CD69+CD103+ cells. (I) Mean proportion of CD4+ T cells expressing 0, 1, 2, 3, 4, 5, or 6 of the cytokines or chemokines IL-17A, TNFα, IFNγ, CCL4, IL-2, or IL-10, categorised by CD69 and CD103 co-expression, from small intestinal biopsies from healthy control subjects (n=10). Statistical analysis performed with one-way ANOVA with Tukey’s multiple comparisons test. *, P ≤ 0.05; **, P ≤ 0.01; ***, P ≤ 0.001.

A donor-derived regulatory CD4+ FOXP3+ population was detected in the scRNAseq data (Fig. 2C), and donor-derived CD25+CD127-CD4+ T cells were detected by flow cytometry in some subjects, consistent with potential long-term residency of SI CD4+ regulatory T cells (Supplementary Fig. 3C). Low cell number of this population precluded further analysis.

CD69+ CD4+ SI T cells showed higher production of multiple cytokines upon short-term stimulation compared with CD69-T cells (Fig. 5C-I). In particular, IL-17A production was almost exclusively restricted to CD69+ T cells (Fig. 5C). CD103+ CD4+ T cells demonstrated higher production of TNFα, CCL4, IL-17A, and IL-10 than CD69+ CD103-counterparts. There was a similar gradient of functionality between the three CD4+ populations, with CD103+ cells showed greatest polyfunctional cytokine production (Fig. 5I), as seen in CD8+ T cells. In summary, CD4+ SI T cell populations demonstrated analogous differences in phenotype and functionality to their CD8+ counterparts.

## Discussion

This study has identified two transcriptionally distinct states of functionally-resident *bona fide* human intestinal CD8+ T_RM_ cells, which differ in phenotype and cytokine production. This was demonstrated using the rigorous approach of identifying donor-derived populations in the intestinal graft post-transplant, which confirms functional tissue residency. These data were replicated both within graft-infiltrating T cell populations establishing residency *de novo*, and in the healthy gut. The CD8+ T_RM_ subsets differ in the expression of several genes previously associated with the human T_RM_ signature, which suggests that these signatures, derived from bulk RNAseq data, may represent an amalgam of transcriptionally distinct T_RM_ subsets (Kumar et al., 2017).

The two identified CD8+ T_RM_ populations differ in IL7R (CD127) expression, and in IL-2 production, which may indicate differing persistence and proliferation properties, as well as in the expression of chemokine receptors and integrins, indicating potential differences in tissue homing and retention. Of particular interest, β2-integrin is more highly expressed on CD69+CD103-T_RM_ cells, a population which remains difficult to positively distinguish from recent immigrants from the circulation. β2-integrin could, in addition to CD69, represent an important positive marker to identify this population.

β2-integrin forms part of the heterodimer LFA-1, which facilitates firm adhesion on blood vessel endothelium via ICAM-1, a critical stage in lymphocyte trafficking (Fagerholm et al., 2019). β2-integrin also regulates the immunological synapse providing a co-stimulatory signal, both in the interaction with antigen-presenting cells, and with infected target cells (Liu et al., 2009). LFA-1 is up-regulated on liver-resident T cells and facilitates their patrolling of hepatic sinusoids, indicating a role for LFA-1 in T_RM_ motility and tissue surveillance (McNamara et al., 2017). Intriguingly, the interaction of LFA-1 with ICAM-1 reduces cellular responses to TGFβ, a key cytokine in T_RM_ development (Verma et al., 2012), while TGFβ can inhibit LFA-1 expression and function (Boutet et al., 2016). This suggests a role for β2-integrin in the development of CD103-CD8+ T_RM_ cells, which may be TGFβ-independent, as in mice (Bergsbaken and Bevan, 2015).

The *ITGB2* locus is a differentially methylated region in the blood and mucosa in IBD, and mucosal gene expression is associated with disease activity (Kennedy et al., 2016; Harris et al., 2014; Román et al., 2013). We hypothesize that this association with IBD, in which T_RM_ cells play a key role (Zundler et al., 2019), could be mediated via altered β2-integrin expression and its effects on T_RM_ cells, or suggest a pathogenic role for one of the two T_RM_ subsets.

Single-cell transcriptional heterogeneity has previously been examined in lung T_RM_ cells, with two potential subsets identified, characterised by predominantly different gene signatures to those in our study (Snyder et al., 2019). However, their putative subsets contained both CD4+ and CD8+ cells co-clustering, which separated clearly in our study. T_RM_ cell phenotype and behaviour in lung and intestine differ substantially, and this finding may be driven by such tissue-specific differences, or be an effect of the higher cell number in our study (Thome et al., 2014). It remains unclear if analogous subsets to the intestinal populations described in this study exist in other tissues, whether such subsets differ in residency characteristics and functional capacity, and whether they play distinct roles in human health and disease.

In conclusion, we have used the human model of intestinal transplantation to study single-cell heterogeneity in the donor-derived *bona fide* T_RM_ intestinal T cells. We found that CD8+ T_RM_ cells form two transcriptionally, phenotypically, and functionally distinct subsets, with parallel findings in CD4+ T cells. In particular, we report the association between β2-integrin and CD69+CD103-intestinal T_RM_ cells, which may prove not only a useful marker for this population, but could also have a role in T_RM_ development and function.

## Materials and methods

### Human biological material

Intestinal transplant recipients were identified via the Oxford University Hospitals NHS Foundation Trust (OUHFT) transplantation service (Oxford, United Kingdom). Healthy control study subjects were identified via the OUHFT endoscopy service at the time of routine endoscopy. Peripheral blood and intestinal biopsies were taken at the time of endoscopy under the study framework and consent of the Oxford Gastrointestinal Illnesses Biobank (REC Ref: 16/YH/0247). Patient and sample characteristics are described in Supplementary Table 1.

Small intestinal biopsies were collected at the time of endoscopy and transported in R10 (RPMI-1640 [Lonza] + 10% FCS [Sigma-Aldrich] + 1% penicillin/streptomycin [Sigma-Aldrich]), before cryopreservation in freezing medium (10% DMSO [Sigma-Aldrich], 90% FCS [Sigma-Aldrich]). This method preserves immune cell viability, surface marker expression, and function (Konnikova et al., 2018). When required, samples were rapidly thawed in a 37°C water bath, washed in 20ml R10 before tissue dissociation.

Duodenal samples were incubated in R10 media with 1mg/ml Collagenase D (Roche) and 100μg/ml DNase (Thermo Fisher Scientific) for one hour in a shaking incubator at 37°C. Biopsies were then dissociated by vigorous agitation using a GentleMACS Dissociator (Miltenyi Biotec), then strained through a 70μm filter. Cells were washed with R10 media. For samples undergoing *ex vivo* stimulation or cell sorting, the mononuclear cells were isolated on a discontinuous 70% and 35% Percoll gradient (GE Healthcare) by centrifugation at 700g for 20 minutes without brake. Mononuclear cells were collected from the interface and washed in R10.

For multiplex fluorescence chip cytometry, single intestinal biopsies were embedded in OCT cryo-embedding matrix (Thermo Fisher Scientific) then frozen in isopentane (Sigma-Aldrich) suspended over liquid nitrogen, and stored at −80°C until use. Peripheral blood mononuclear cells (PBMCs) were isolated from subject blood samples by density gradient centrifugation. In brief, blood was diluted 1:1 with PBS, then layered onto Lymphoprep (Axis-Shield), and centrifuged at 973g for 30 minutes without brake. The mononuclear layer was collected and washed with R10. Any remaining red blood cells were lysed with ACK (Ammonium-Chloride-Potassium) solution for 2-3 minutes, and washed again in R10, before cryopreservation in freezing medium as above.

### *Ex vivo* stimulation

*Ex vivo* stimulation was performed as previously described (Provine et al., 2018). In brief, purified intestine-derived mononuclear cells were plated at approximately 1×10^6^/well in a U-bottom 96-well plate. Cell stimulation cocktail containing phorbol 12-myristate 13-acetate (PMA) and ionomycin (Biolegend) was added in accordance with manufacturer’s instructions in the presence of brefeldin A and monensin (both Biolegend). Cells were incubated at 37°C, 5% CO_2_, for 4 hours.

### Flow cytometry and cell sorting

For surface marker staining, cells were stained in 50μL of FACS buffer (PBS + 1mM EDTA + 0.05% BSA) for 30 minutes at 4°C. Surface antibodies and clones used were: CD3 (OKT3 or UCHT1), Vδ2 (B6), CD4 (OKT4 or SK3), Vα7.2 (3C10), CD38 (HIT2), CD103 (Ber-ACT8), γδTCR (11F2), CD69 (FN50), HLA-A2 (BB7.2), HLA-A3 (GAP.A3), CD8 (SK1 or RPA-T8), CD161 (191B8), nearIR and zombie yellow fixable viability dye (Thermo Fisher Scientific or Biolegend), β2 integrin/CD18 (L130), CD25 (2A3), KLRG1 (REA261), CD127 (A019D5), CD7 (M-T701).

For intracellular cytokine staining, after surface staining as above, cells were fixed and permeabilised in 100μL Cytofix/Cytoperm solution for 20 minutes at 4°C (BD Biosciences). Cells were then washed twice in Perm/Wash buffer (BD Biosciences). Intracellular staining was performed in 50μl of Perm/Wash buffer (BD Biosciences) for 30 minutes at 4°C using the following antibodies and clones: CCL4 (D21-1351), Ki67 (B56), TNFα (MAb11, IL-17A (BL168), IFNγ (B27), IL-2 (MQ1-17H12), GzmK (GM26E7), IL-10 (JES3-9D7).

After staining, cells were stored at 4°C protected from light until data acquisition. Flow cytometry data were acquired on a BD LSRII flow cytometer (BD BioSciences) or Aurora spectral flow cytometer (Cytek). For fluorescence-activated cell sorting (FACS) samples were surface stained as above, with DAPI (Thermo Fisher Scientific) used as viability dye. FACS was performed on an AriaIII (BD Biosciences; 70μm nozzle). Antibodies were purchased from BioLegend, BD Biosciences, Miltenyi Biotec, or Thermo Fisher Scientific.

### Flow cytometry data analysis and statistics

Flow cytometry data were analysed using FlowJo version 9.9.5 and version 10.6.1 (FlowJo, LLC). Statistical analyses of flow cytometric data were performed using Prism version 8 (Graphpad Software). For phenotypic analysis of rare cell subsets, populations with fewer than 10 cells were excluded.

### Chip cytometry

Samples were frozen in OCT (as described above), cryosectioned onto coverslips and placed in cytometer chips (Zellsafe Tissue chips, Zellkraftwerk, GmbH, Deutscher Platz 5c, 04103 Leipzig, Germany). Sections were fixed *in situ* at room temperature for 10 minutes using 4% paraformaldehyde, then washed with 10 mL of PBS. Non-specific binding was blocked by incubating in 5% goat serum (Thermo Fisher Scientific) in PBS for 1 hour at room temperature. Fluorophore-conjugated antibodies (as above, plus pan-cytokeratin (C-11, GeneTex), and Histone-H3 (17H2L9, Thermo Fisher Scientific)) were diluted for staining in PBS. Immunostaining was performed iteratively, with up to three colours applied simultaneously. Fluorophores were then bleached and additional antibodies applied to build up the panel (Hennig et al., 2009). Images were acquired using a Zellscanner One Chip cytometer (Zellkraftwerk) and ZellExplorer software.

### 10x Genomics library preparation and sequencing

scRNAseq libraries were generated using 10x Genomics Chromium Single Cell V(D)J Reagents Kits (v1 Chemistry) following manufacturer’s instructions. Cells were resuspended in PBS with 0.04% BSA at ∼1000 cells/μL and loaded onto a single lane of the Chromium Controller. Captured cell number was 1,774. Library quality and concentration was determined using a TapeStation (Agilent) and Qubit 2.0 Fluorometer (Thermo Fisher Scientific). Libraries were sequenced on an Illumina HiSeq 4000 as per manufacturer’s instructions to a mean depth of 63,471 mean reads/cell. Library generation and sequencing were performed at the Oxford Genomics Centre (Wellcome Centre for Human Genetics, University of Oxford).

### Smart-Seq2 library preparation and sequencing

Single cells were index-sorted into 96-well plates with one cell per well in 2.3 μl lysis buffer (0.8% (vol/vol) Triton X-100 and 2 U/μL RNase inhibitor). Smart-Seq2 libraries were generated following the published protocol with External RNA Controls Consortium (ERCC) RNA (1:100,000) added prior to sequencing (Picelli et al., 2014). cDNA was pre-amplified by PCR (21 cycles). Libraries were sequenced on an Illumina HiSeq4000 with 75bp paired-end reads. Library generation and sequencing were performed at the Oxford Genomics Centre (Wellcome Centre for Human Genetics, University of Oxford).

### Droplet-based (10x Genomics) scRNAseq data analysis

FastQ generation, read alignment, barcode counting and UMI counting were performed using the Cell Ranger pipeline v2.2.0. Downstream processing steps were performed using Seurat v2.3.4 (Butler et al., 2018; Stuart et al., 2019). Briefly, TCR and BCR genes, and genes expressed in fewer than 10 cells, were removed. Cells with < 3460 UMIs (local minimum of the UMI distribution to the left of the mode UMI count), < 500 genes, and > 10,000 UMIs, > 2500 genes, and/or > 10% mitochondrial reads were removed (Supplementary Fig. 2A). Variable genes were identified using M3Drop (Andrews et al., 2018). Data were log normalised and scaled, with cell-cell variation due to UMI counts, percent mitochondrial reads, and S and G2M cell cycle scores regressed out. The top 10 principal components (PCs) were used as input for graph-based clustering (0.4 resolution), as determined by visual inspection of the scree plot. Clusters were visualised by tSNE. Differential gene expression analysis between clusters was performed using the Wilcoxon Rank Sum test (FindMarkers function with default parameters).

### Plate-based (Smart-Seq2) scRNAseq data analysis

Reads were trimmed to remove contaminating adapter and oligo-dT primer sequences using Trimmomatic v0.36 (Bolger et al., 2014). Trimmed reads were aligned to the human genome (hg38 assembly) plus added ERCC “spike-in” sequences using STAR v2.5.3a (--outFilterMismatchNoverLmax 0.04 --outFilterType BySJout -- outMultimapperOrder Random) (Dobin et al., 2012). Alignments were filtered using Samtools v1.6 (Li et al., 2009) to retain only primary alignments and properly paired reads. Ensembl gene counts were generated using featureCounts v1.6.0 (-C -B -p) (Liao et al., 2014). Poor quality cells that fit one or more of the following criteria were removed from the analysis (Supplementary Fig. 2G-I): small log-library size (< 3 median absolute deviations (MADs) below the median), low percentage of uniquely mapped reads (< 55%), low gene count (< 3 MADs below the median), high percentage of ERCCs (> 37.5%), or high mitochondrial read fraction (> 6%). Outlier cells with a large library size or high gene count (potential doublets) were also removed. Genes with “undetectable” expression were removed (gene defined as “detectable” if at least 5 read counts in two cells), along with TCR (and BCR) genes. Log normalised expression values were generated using the normalize function from scater v1.10.1 (McCarthy et al., 2017) with cell-specific size factors calculated using scran v1.12.1 (Lun et al., 2016). Feature selection was performed using M3Drop v3.10.4 (FDR < 0.01) (Andrews et al., 2018) and the expression matrix subset to retain only the selected genes. Clustering analysis was performed using SC3 v1.12.0 (Kiselev et al., 2017). Clusters were visualised by UMAP generated using the top 30 PCs (McInnes et al., 2018) (generated using the top 30 principal components). Differential gene expression analysis was performed using DEsingle v1.2.1 (Miao et al., 2018) (FDR < 0.05).

Upon publication, data will be deposited in the NIH GEO database.

## Supplemental material

- Supplemental Figure 1
- Supplemental Figure 2
- Supplemental Figure 3
- Table 1: Subject and sample characteristics
- Table 2: Differentially expressed gene list

## Author contribution

MF, NP, LG, PK, and PA conceived and designed the study. Patient identification, consent, and sample collection was performed by MF, TA, PF, GV, SR, and PA. Experiments and analyses were performed by MF, NP, LG, KP, AA, SI, and HF. Cell sorting was performed by HF. MF, NP, and LG created the figures. MF drafted the article. MF, NP, LG, KP, AA, SI, HF, TA, PF, GV, SR, ES, PK, and PA critically revised the article and approved the final version.

## Acknowledgments

We thank the Oxford TGU GI illnesses biobank team, and the clinicians and endoscopists at Oxford University Hospitals NHS Foundation Trust, for assistance in patient identification and sample collection. We are grateful to Dr Chris Willberg and Joachim Hagel at the Peter Medawar Building for work establishing the chip cytometry facility. We are grateful to Professor Laura Mackay for her advice in the conception of this project.

We thank the Oxford Genomics Centre at the Wellcome Centre for Human Genetics (funded by Wellcome Trust grant reference 203141/Z/16/Z) for the generation of sequencing data. The research was funded by the Wellcome Trust (WT 109665MA), NIHR Senior Fellowship (PK), and National Institute for Health Research (NIHR) Oxford Biomedical Research Centre (BRC). The authors have no additional financial interests. The views expressed are those of the author(s) and not necessarily those of the NHS, the NIHR or the Department of Health.

## Supplemental Figure 1

(A) Illustration demonstrating the principle of HLA allele congenic cell tracking to identify donor- and recipient-derived T cell populations in the transplanted small intestinal graft using fluorophore-conjugated antibodies to Class I HLA haplotypes discordant between donor and recipient.

(B) Representative flow cytometry plot of Class I HLA non-specific staining (0.01-0.12%) from a non-transplant PBMC sample. Gated on Live CD3+ T cells.

(C) Representative flow cytometry gating scheme for identification of T cell subsets.

(D) Representative histogram of CD161 expression of paired donor- and recipient-derived CD8+ intestinal T cells.

(E) Percentage of CD161+ donor- and recipient-derived CD8+ intestinal T cells categorised by time post-transplant (left), or grouped, with paired samples connected by black lines (right) (n=15; mean +/- SEM).

Statistical analysis performed with Wilcoxon matched-pairs signed rank test. *, P ≤ 0.05.

## Supplemental Figure 2

(A) Quality control parameters and thresholds for 10x Genomics scRNAseq Experiment 1. Number of genes per cell (left), number of unique molecular identifiers (UMIs) per cell (centre), and percentage of mitochondrial reads per cell (right) are displayed. Each dot represents a single cell, with red and blue lines indicating maximum and minimum thresholds, respectively.

(B) Dot plot showing expression in Experiment 1 of *ITGB2* and its potential heterodimeric partner, *ITGAL* in Experiment 1. Other potential heterodimeric alpha integrin partners, *ITGAD*, *ITGAM*, and *ITGAX* were not detected. Dot size indicates the proportion of cells in which the gene is present. Colour intensity indicates the mean expression level of the gene.

(C) Violin plots showing expression of *ITGB2* and its potential heterodimeric partner *ITGAL* in Experiment 1. Other potential heterodimeric alpha integrin partners, *ITGAD*, *ITGAM*, and *ITGAX* were not detected.

(D) Dot plot showing expression in Experiment 1 of 11 genes previously negatively associated with tissue residency (downregulated in human CD69+ cells in comparison to CD69-T cells; (Kumar et al., 2017)). Dot size indicates the proportion of cells in which the gene is present. Colour intensity indicates the mean expression level of the gene. Other genes in the geneset were not detected in the data (*SBK1*, *NPDC1*, *KRT72*, *SOX13*, *KRT73*, *TSPAN18*, *PTGDS*).

(E) Violin plots showing expression in Experiment 1 of 11 genes previously negatively associated with tissue residency (downregulated in human CD69+ cells in comparison to CD69-T cells; (Kumar et al., 2017)).

(F) Violin plots showing expression in Experiment 1 of 13 genes previously associated with tissue residency in human CD69+ T cells (Kumar et al., 2017), demonstrating discordant expression in conventional T cell clusters.

(G-I) Quality control parameters and thresholds for Smart-Seq2 scRNAseq Experiment 2.

(G) Histogram of number of reads per cell and (H) number of genes per cell. Red and blue lines indicate maximum and minimum thresholds, respectively.

(I) Plot showing the percentage of External RNA Controls Consortium (ERCC) spike-ins per cell against the number of genes per cell. Black line indicates upper threshold.

(J) Violin plots showing expression of *ITGB2* and its potential heterodimeric partners

*ITGAL* and *ITGAX* in Experiment 2. *ITGAD* and *ITGAM* were not detected in the data.

## Supplemental Figure 3

(A) Representative expression of CD161, CD7, CD127 (IL7R), ITGB2, and Granzyme K on CD103-(red) and CD103+ (black) donor-derived T cells from the intestinal transplant graft.

(B) The proportion of CD8+ T cells co-expressing CD69 and CD103 in small intestinal biopsies from the duodenum (duo) or ileum from healthy control subjects (n=5 for both duodenum and ileum). Blue, CD69-103-cells; Red, CD69+CD103-cells; Black, CD69+CD103+ cells.

(C) Representative flow cytometry gating of CD4+ CD25+CD127lo T cells.

Percentage of donor- or recipient-derived intestinal CD4+ CD25+CD127lo T cells. Connecting lines represent populations from the same subject (n=23).

