## Supplementary figures and images for "Human intestinal tissue-resident memory CD8+ T cells comprise transcriptionally and functionally distinct subsets"

### Supplemental Figure 1

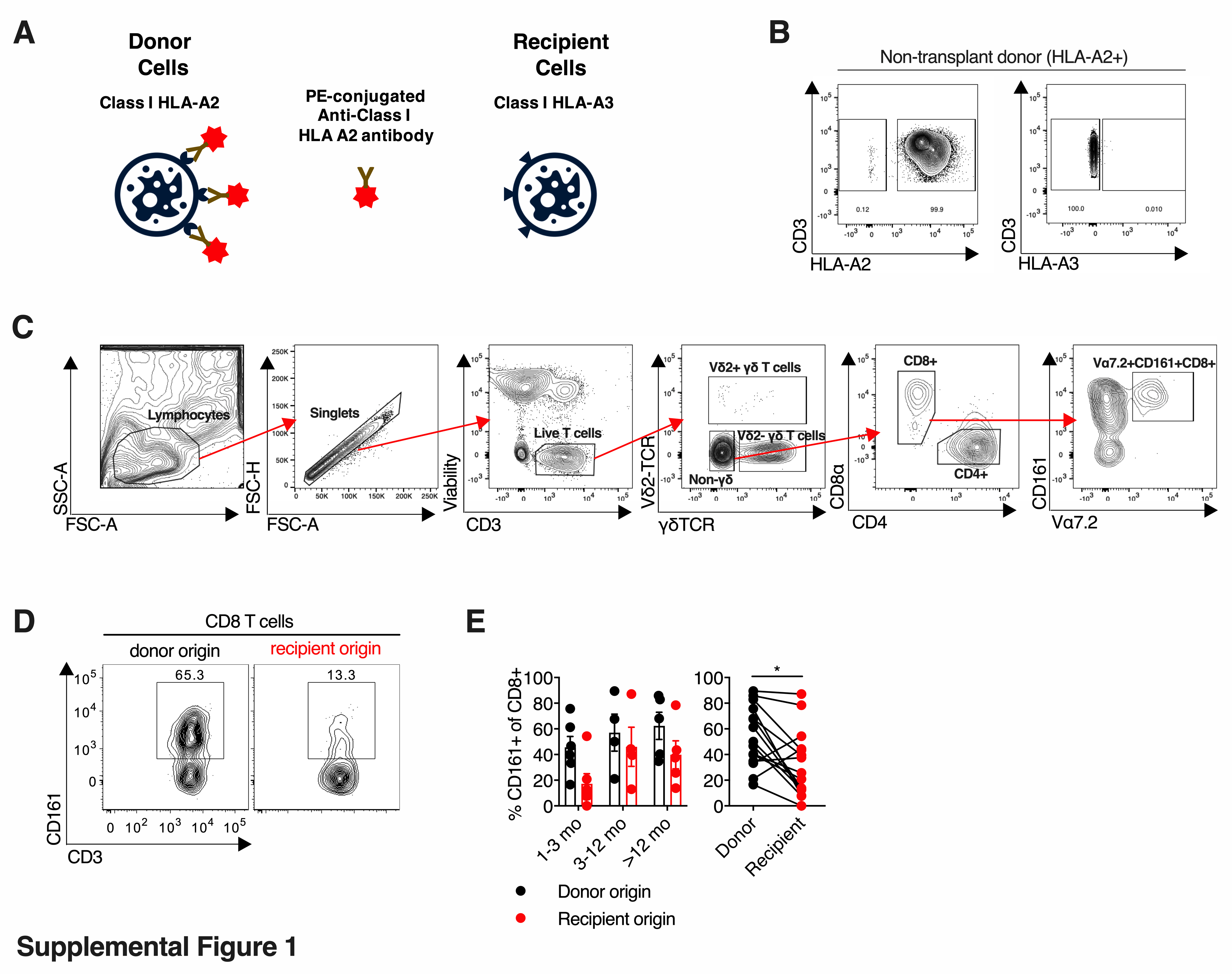

### Supplemental Figure 2

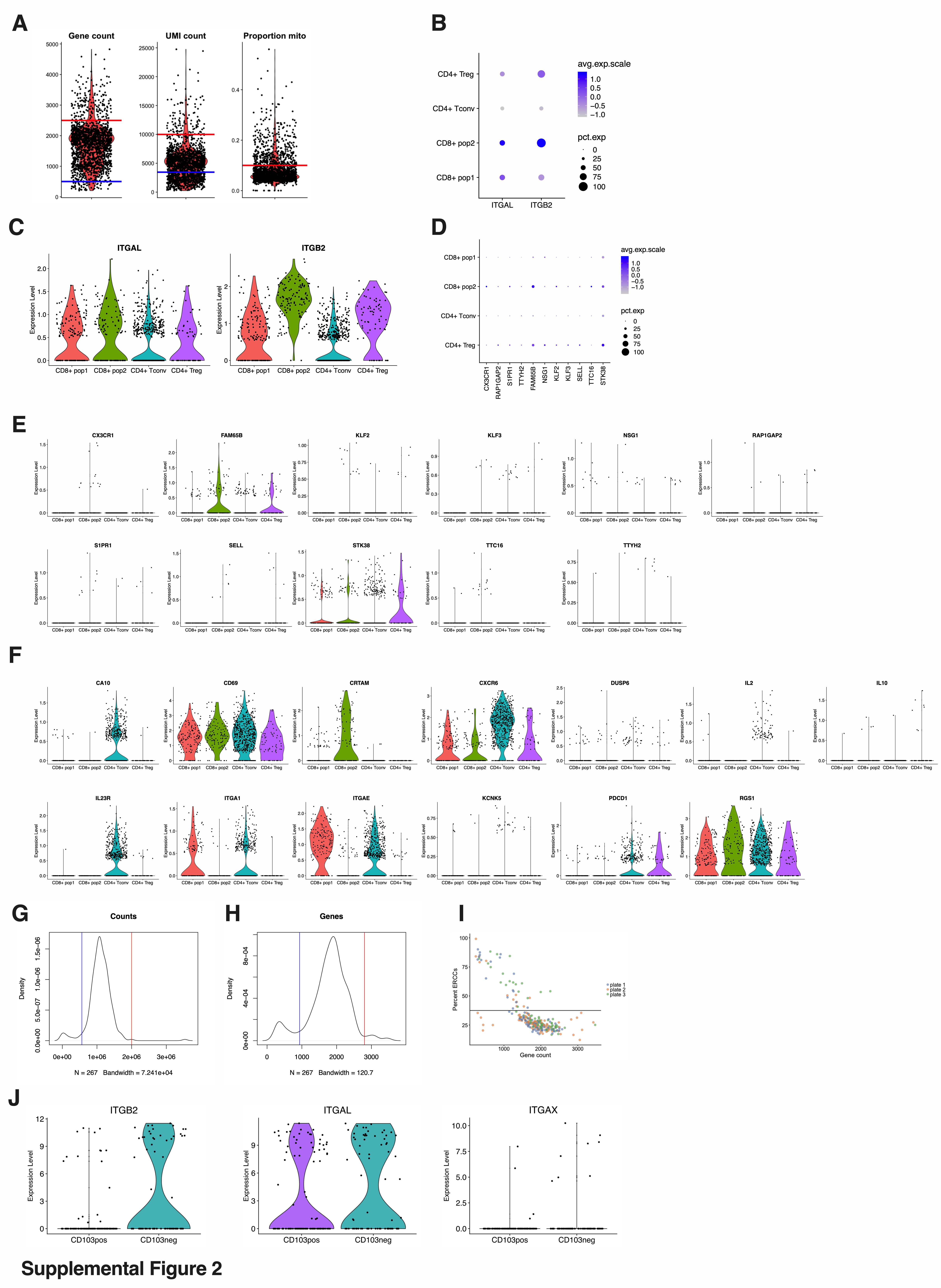

### Supplemental Figure 3

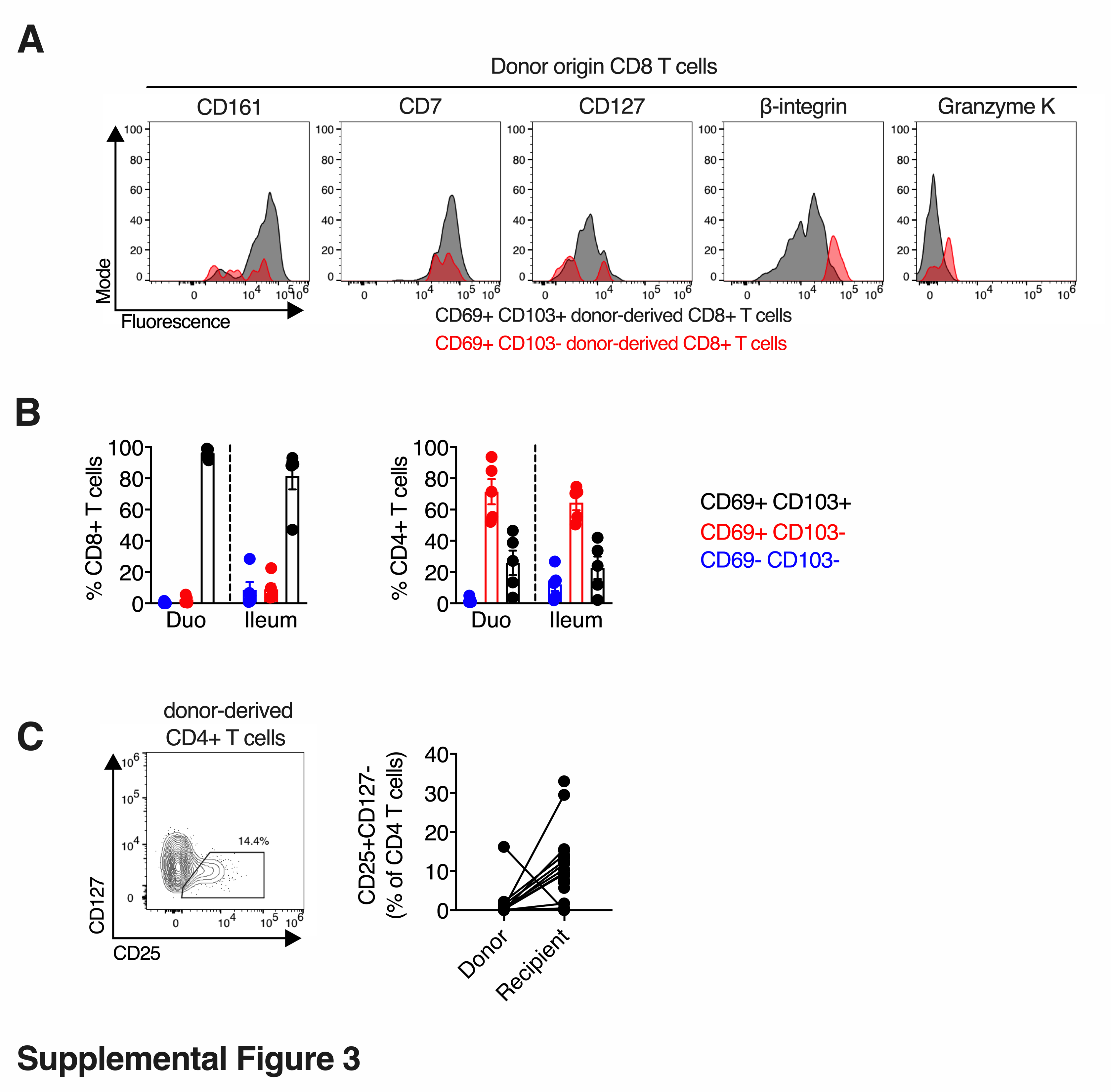
